# TNER: A Novel Background Error Suppression Method for Mutation Detection in Circulating Tumor DNA

**DOI:** 10.1101/214379

**Authors:** Shibing Deng, Maruja Lira, Stephen Huang, Kai Wang, Crystal Valdez, Jennifer Kinong, Paul A Rejto, Jadwiga Bienkowska, James Hardwick, Tao Xie

**Affiliations:** Pfizer Early Clinical Development Biostatistics, 10770 Science Center Drive San Diego, CA, USA; Pfizer Early Oncology Development and Clinical Research, 10770 Science Center Drive San Diego, CA, USA

## Abstract

The use of ultra-deep, next generation sequencing of circulating tumor DNA (ctDNA) holds great promise for early detection of cancer as well as a tool for monitoring disease progression and therapeutic responses. However, the low abundance of ctDNA in the bloodstream coupled with technical errors introduced during library construction and sequencing complicates mutation detection. To achieve high accuracy of variant calling via better distinguishing low frequency ctDNA mutations from background errors, we introduce TNER (Tri-Nucleotide Error Reducer), a novel background error suppression method that provides a robust estimation of background noise to reduce sequencing errors. It significantly enhances the specificity for downstream ctDNA mutation detection without sacrificing sensitivity. Results on both simulated and real healthy subjects’ data demonstrate that the proposed algorithm consistently outperforms a current, state of the art, position-specific error polishing model, particularly when the sample size of healthy subjects is small. TNER is publicly available at https://github.com/ctDNA/TNER.

## 1. Introduction

Cancer is a genetic disease that is driven by changes to genes controlling cellular function (Hanahan and Weinberg, 2011). Characterizing the disease at the molecular level is essential to early detection, personalized therapy based on tumor genomic profile, monitoring tumor progression and response to treatment as well as identification of resistant mechanisms (Diehl, et al., 2008). This typically requires tumor tissue biopsies to obtain samples for genotyping or other molecular analyses for solid tumors. Biopsy procedures are usually invasive, and come with additional risk to patient’s health. In many cases, tumor tissue biopsy is contraindicated medically and the tissue samples are often insufficient or unsuitable for molecular profiling (Kinde, et al., 2011). In addition, cancer is a heterogeneous disease with different subclones within the same primary tumor and between the primary tumor and metastatic lesions. This heterogeneity in tumor can lead to variations in tumor tissue sampling through biopsy (Venesio, et al., 2017).

Both cancer and normal cells shed DNA as a result of apoptosis and other biological processes, and release DNA fragments into the blood stream to become cell-free DNA (cfDNA)(Jiang, et al., 2015; Snyder, et al., 2016; Underhill, et al., 2016). cfDNA derived from tumor cells is called circulating tumor DNA (ctDNA), and provides a real time genomic snapshot of cancer cells due to the relatively short half-life of cfDNA (~1-2 hours)(Diehl, et al., 2008; Volik, et al., 2016). Thus ctDNA is a form of “liquid biopsy” that provides a non-invasive alternative to tissue biopsy for cancer diagnosis and monitoring (Bettegowda, et al., 2014; Lippman and Osborne, 2013). Moreover, ctDNA generally comes from all tumor lesions and is pooled in the circulatory system, therefore it can reduce sampling variation associated with tumor heterogeneity in comparison to a single tissue biopsy (Crowley, et al., 2013).

The fraction of ctDNA in the total cfDNA in plasma, however, can be extremely low in many cancer patients (Diehl, et al., 2008; Volik, et al., 2016). Recently established techniques such as droplet-digital PCR (ddPCR) enable detection and quantification of low abundance ctDNA, but with limitation to cover only a small number of known “hotspot” mutations (Openshaw, et al., 2016; Volik, et al., 2016). Advances in DNA sequencing technology have made it possible to identify ctDNA mutations at comparable sensitivity to ddPCR (Chen, et al., 2017; Thierry, et al., 2014) when the sequence coverage is sufficient (>10,000× per base). One of the most significant challenges in detecting ctDNA mutations is suppressing technical errors introduced during library preparation, PCR amplification or the sequencing itself (Nakamura, et al., 2011). While errors arising during PCR amplification can be removed effectively using molecular barcodes (Nakamura, et al., 2011), other technical errors are more universal and also need to be removed before mutation calling (Kinde, et al., 2011; Kirsch and Klein, 2012). Newman *et al* (Newman, et al., 2016) recently proposed a creative integrated digital error suppression (iDES) method that includes both a molecular barcoding system to reduce PCR errors and a background polishing model with an improved estimation of background mutation error rate (BMER) compared to the previous computational method used in CAPP-Seq (Newman, et al., 2014). Specifically, the BMER was mostly estimated using a model of Gaussian distribution on the mutation data from a collection of healthy subjects (Newman, et al., 2016). As far as we know, there are very few back-ground polishing methods designed for ctDNA detection and iDES is the only publically available state of the art method. The polishing method used in iDES increased the percentage of error-free positions from ~90% to ~98% (based on a 300kb panel, Fig 2b in Newman, et al., 2016). However, that means still ~6,000 positions containing a substantial number of noisy bases that could be mis-classified, due to the relatively small sample size (n=12) and the nature of the data (small discrete count) which made it difficult for the Gaussian model to robustly estimate the background.

To provide a more robust estimation of background noise and remove the sequencing artifacts more effectively for panel sequencing data, we developed a novel background polishing method called TNER (Tri-Nucleotide Error Reducer) with a Bayesian consideration to overcome the small sample size issue. TNER is based on tri-nucleotide context data and uses a binomial distribution for the mutation error count to estimate the background from healthy subjects. Tri-nucleotide context (TNC hereafter) are 96 distinct substitutions in specific context of tri-nucleotide, consisting of the 6 distinguishable single nucleotide substitutions (C>A, C>G, C>T, A>C, A>G and A>T) and the 16 possible combinations of immediately preceding and following bases. TNC has been extensively studied in cancer genetics to construct mutation signatures as a response to carcinogens (a great summary at: http://cancer.sanger.ac.uk/cosmic/signatures), to compare the mutational spectra of trunk and branch mutations, or to predict the clinical implications of called mutations (Alexandrov, et al., 2013; Gerlinger, et al., 2012; Rosenthal, et al., 2016). Given that the pattern of low frequency technical errors from next-generation sequencing (NGS) should be similar in normal control samples and patient samples, we argue that local sequence context could help better model noise for small sample size of healthy subjects by leveraging information from other bases with a shared TNC. The TNER methodology proposed here, to the best of our knowledge, is novel in this area. As an effective error reducer, TNER can be easily integrated into an existing variant calling pipeline before the variant caller when applied to detect very low frequency mutations in liquid biopsy samples. TNER is freely available at https://github.com/ctDNA/TNER.

## 2. Methods

### 2.1. NGS data for analysis

To demonstrate the performance of the error suppression model in detecting single nucleotide variations, we analyzed targeted sequencing data of plasma cfDNA of healthy subjects using a panel of 87 cancer genes described previously (Lira et al. AACR 2017, #2749). The barcoded target-enriched DNA library (147Kb) was sequenced on an Illumina HiSeq 4000 platform generating ultra-deep coverage with an average coverage per base of ~12,000.

### 2.2. Tri-nucleotide error reduction model

The detection of ctDNA is typically achieved through detecting signature mutations associated with tumors in cfDNA. Sequence data from cfDNA has many stereotypical errors or other background mutation errors that are not of tumor origin (Park, et al., 2017). In order to call a mutation in ctDNA, the distribution of the BMER needs to be characterized at each nucleotide base position to reduce the false positive error, for example, by modeling cfDNA data on the same NGS panel from healthy subjects (Newman, et al., 2016). The mutation rates from healthy subjects are assumed to be background mutation noise associated with both technical and biological sources. One challenge in characterizing the individual nucleotide BMER from healthy subjects is the relatively small cohort size. The iDES method used 12 healthy subjects (Newman, et al., 2016); we used a comparably sized set of 14 healthy subjects. These small sample sizes do not allow building a reliable estimate of the background error distribution for individual nucleotides. Bayesian method with prior information can help to overcome this limitation.

To better estimate the BMER distribution, we propose a background error model originated from a hierarchical Bayesian method that utilizes the distribution of mutation error rate in a TNC, which consists of the mutated nucleotide and the combinations of immediately preceding and following nucleotides. Mutation signatures characterized by TNC have been used frequently in cancer genetics (Alexandrov, et al., 2013; Gerlinger, et al., 2012; Yang, et al., 2015). There are 96 distinct TNC and we assume they are independent. For a nucleotide in TNC group *i*(*i* =1,…, 96) at base position *j* (*j*=1,…*J*), the number of background error reads *X*_ij_ observed for a given coverage *N*_j_ is assumed to follow a binomial distribution

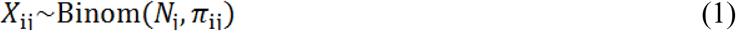

with position specific mutation error rate parameter π_ij_. J is the total number of bases in the panel (147k). With a large *N* (>1,000 typically) and a small π(<1%), X can also be modeled as a Poisson distribution

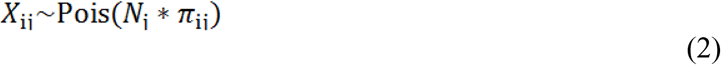

with rate parameter N_j_ * π_ij_. We will focus on the binomial model here.

The BMER at position j can be estimated using the average mutation error rate of the j^th^ base position from the 14 healthy subjects, 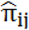. This position specific parameter will be poorly estimated with the small sample size. To improve the estimate of π (for simplicity we drop the subscription for now), we propose a Bayesian framework and assume that π follows a beta distribution within a TNC

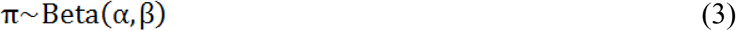

The use of beta prior is primarily due to its conjugation to binomial distribution and also due to its goodness of fit to the data (see discussion). For convenience, we re-parameterize the beta distribution using its mean as a parameter.

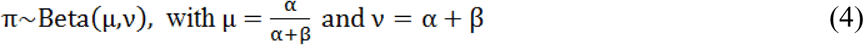

The prior parameters of the beta distribution can be estimated based on the BMER distribution of nucleotides in a TNC using method of moments (Bowman and Shenton, 1998). The mean parameter μ can be estimated by the average mutation error rate 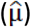 of nucleotides in the TNC. The *ν* parameter can be estimated using 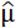 and the sample variance of BMER within the TNC. For a position with *x* mutation count out of *n* total reads, the posterior distribution of the BMER at this position will be a Beta(α + *x*, β + n − *x*) with a mean parameter

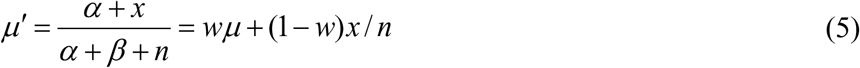

where w=(α+β)/(α+β+n).

Therefore, the posterior mean of the position specific BMER for position *j* with TNC *i* can be estimated with a shrinkage estimator, that is, a weighted average of the TNC level mutation error rate 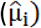 and the position specific rate 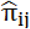

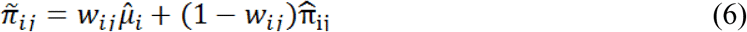

The weight *w*_ij_ can be derived in closed form under beta-binomial distribution and estimated using method of moments (Colews, 2013). We found the analytic Bayesian weight worked well for vast majority of the positions except for a small number of (<1%) positions where the estimated position specific error rate 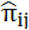 is large. In those positions the shrinkage towards a smaller 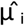 tends to underestimate the true background mutation error. Therefore we adopted a modified weight that balances the relative size of the TNC error rate and the position specific error rate

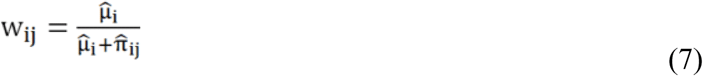

This weight function provides less shrinkage when the position specific mutation error rate is high - a property that helps retain the position specific background when it is much higher than the tri-nucleotide level background. Although this simple weight does not reflect the impact of sample size, a larger sample size helps provide a better estimate of π_ij_. Due to this modification in weight, TNER adopted a more heuristic approach than a full Bayesian method.

Once we have an estimate of the BMER π_ij_ using eq. (6), the threshold for mutation detection can be defined based on the upper posterior credible interval bound of π_ij_. At -α level, the upper 1-α/2 Clopper-Pearson interval bound for a binomial proportion is

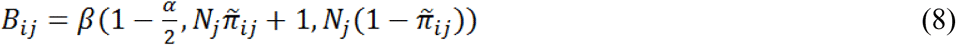

where β() is the quantile function of beta distribution; 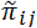 is the posterior estimate of mutation error rate in Eq (6) and N_j_, is the average total reads for this position from healthy subjects. If the observed mutation error rate at position *j* with TNC *i* is lower than *B_ij_*, those variants mapped to the TNC will be classified as background noise and polished using the reference allele, and otherwise the variants will not be polished (possibly true mutations). In the Bayesian model, multiple comparison is not a major concern as the prior distribution allows pooling information between positions and avoids false positive call when variation is low (Gelman, et al., 2012). In our analysis, the false positive calls are very rare when applying the method to the healthy subjects (see Results). Similar beta-binomial model has been used in other studies (Gerstung, et al., 2014; He, et al., 2015; Martincorena, et al., 2015). However, none of them used the model to estimate the BMER distribution with TNC, nor did they apply to ctDNA NGS data.

## 3. Results

We first evaluated the TNER model on the healthy subject data using the leave-one-out method. We built the background model using data from 13 healthy subjects and predicted the mutation in the leave-out subject. Similar to Newman et al (Newman, et al., 2016), we counted the number of error free positions, defined as those positions with exclusively reference allele reads after error suppression, for each of the 14 healthy subjects at all 147k nucleotide positions, and compared the different error suppression methods, including background polishing from iDES and the TNER method (Figure 1). For TNER we used α=0.01, although the results were similar for α=0.05. We also calculated the panel-wide error rate which is defined as the number of non-reference allele reads (frequency<5%, to exclude SNPs) divided by the total reads. TNER method has the highest error-free positions and lowest panel-wide error rate, demonstrates its superior specificity in reducing false positive error.

**Figure 1.**
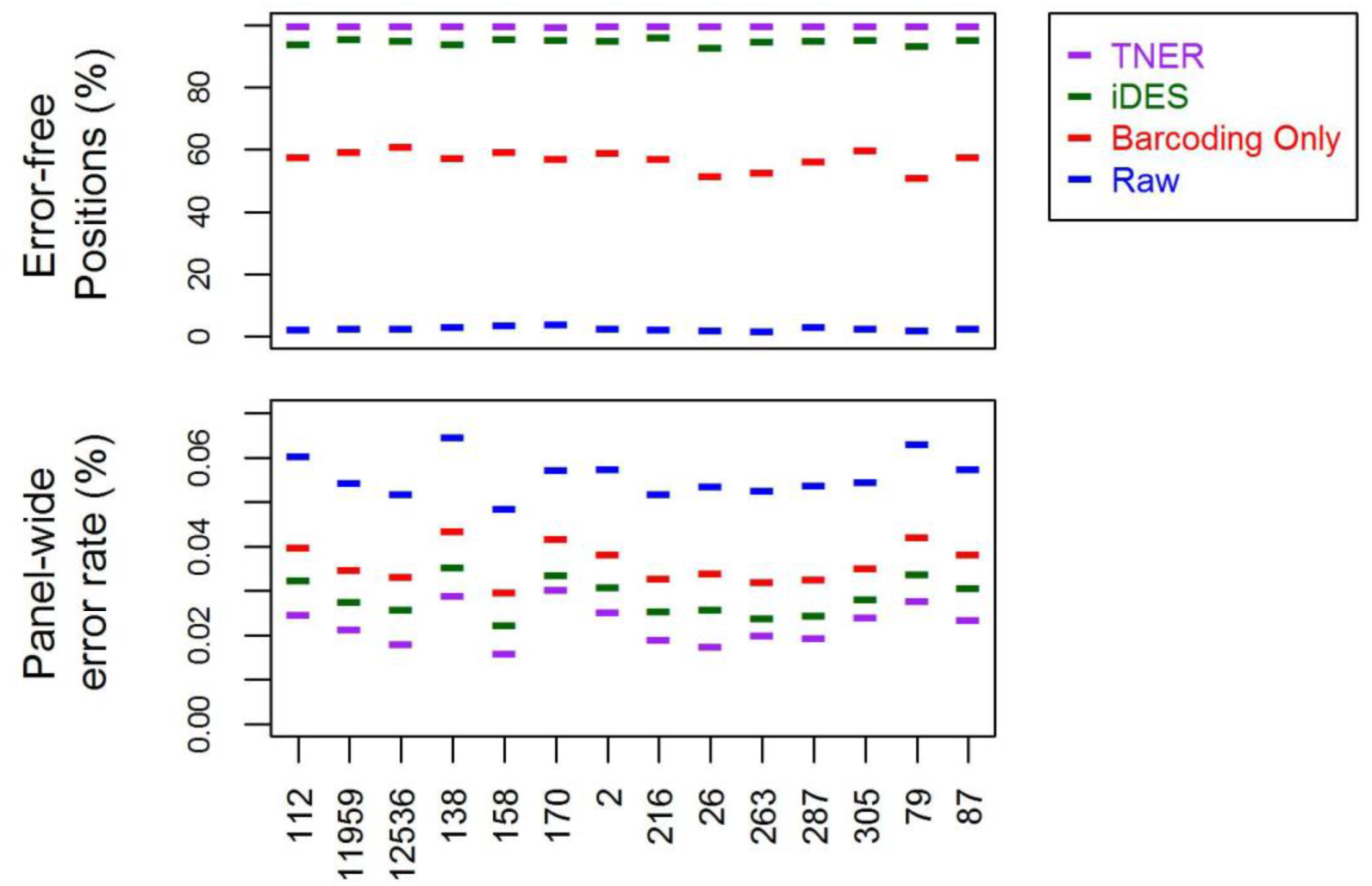
Error free position (%) and panel-wide error rate of the 14 healthy subjects’ data (sample labels on x-axis) from the leave-one-out analysis with different methods. Raw = raw data, Barcoding Only = Barcoding error reduction only.

To test the sensitivity of the method, we used data from three healthy subjects that were not part of the background cohort. One subject had 10 unique private SNPs that were not shared by any of the healthy subjects. We did an in-silico experiment to dilute this subject’s data with the other two health subjects in a 1:250:250 ratio and assume heterozygosity so we have an expected allele frequency of 0.1% for the 10 private SNPs. We found both iDES and TNER (α=0.01) were able to detect all the 10 SNPs in this experiment.

To compare the performance of the position specific background polishing method and the TNER method more rigorously, we evaluated their sensitivity and specificity at various detection thresholds using simulation studies (see the schematic in Supplementary Figure 1). The simulation used the average position specific mutation error rate from the 14 healthy subjects as the BMER which is a matrix of 147k rows and four columns. Each column is a nucleotide that the reference base can mutate to, including the reference nucleotide which is zero. We randomly selected 1,000 bases (rows) out of the 147k total, and then at each of the selected base a simulated allele frequency (simulated signal) was added to the existing BMER of a selected non-reference nucleotide (column). Specifically, for each of the 1,000 positions, there are three possible non-reference nucleotides it can mutate to. We chose the nucleotide with the largest BMER value as the selected nucleotide to add the simulated signal. If the BMER are all zeros at this position, we used the first non-reference letter (A-C-T-G) as the selected nucleotide to add signal. This updated BMER matrix is the same as the original matrix except that 1,000 rows have a signal added to a selected column; with the updated BMER matrix we simulated the read counts with a total coverage of 10,000 per position using a binomial and a normal distribution. For a normal distribution, we simulated the allele fractions with the updated BMER as mean and the square root of the BMER divided by 100 as standard deviation. The read counts are calculated by multiplying the simulated allele fractions with the total coverage of 10000 (round to whole number). The simulated counts were further split into forward and reverse strand with a random forward to reverse strand ratio centered around 1. The TNER method and the position specific Gaussian models from the iDES were then separately applied to the simulated data. As the true positives and true negatives are known, sensitivity and specificity were calculated under various detection thresholds (α values). Receiver operating characteristic (ROC) curves in Figure 2 compare the two methods under different scenarios. The TNER method performed better than the position specific Gaussian model in all cases – data simulated under different distributions and different mutation rate (MR) as shown by ROC curves. The simulated mutation signal 0.075% and 0.1% were chosen because they are close to the limit of detection for the methods when per base coverage is around 10,000x. Larger signals will be too easy to detect by either methods and smaller signals will be lower than the detection limit and difficult to be detected by any methods.

**Figure 2.**
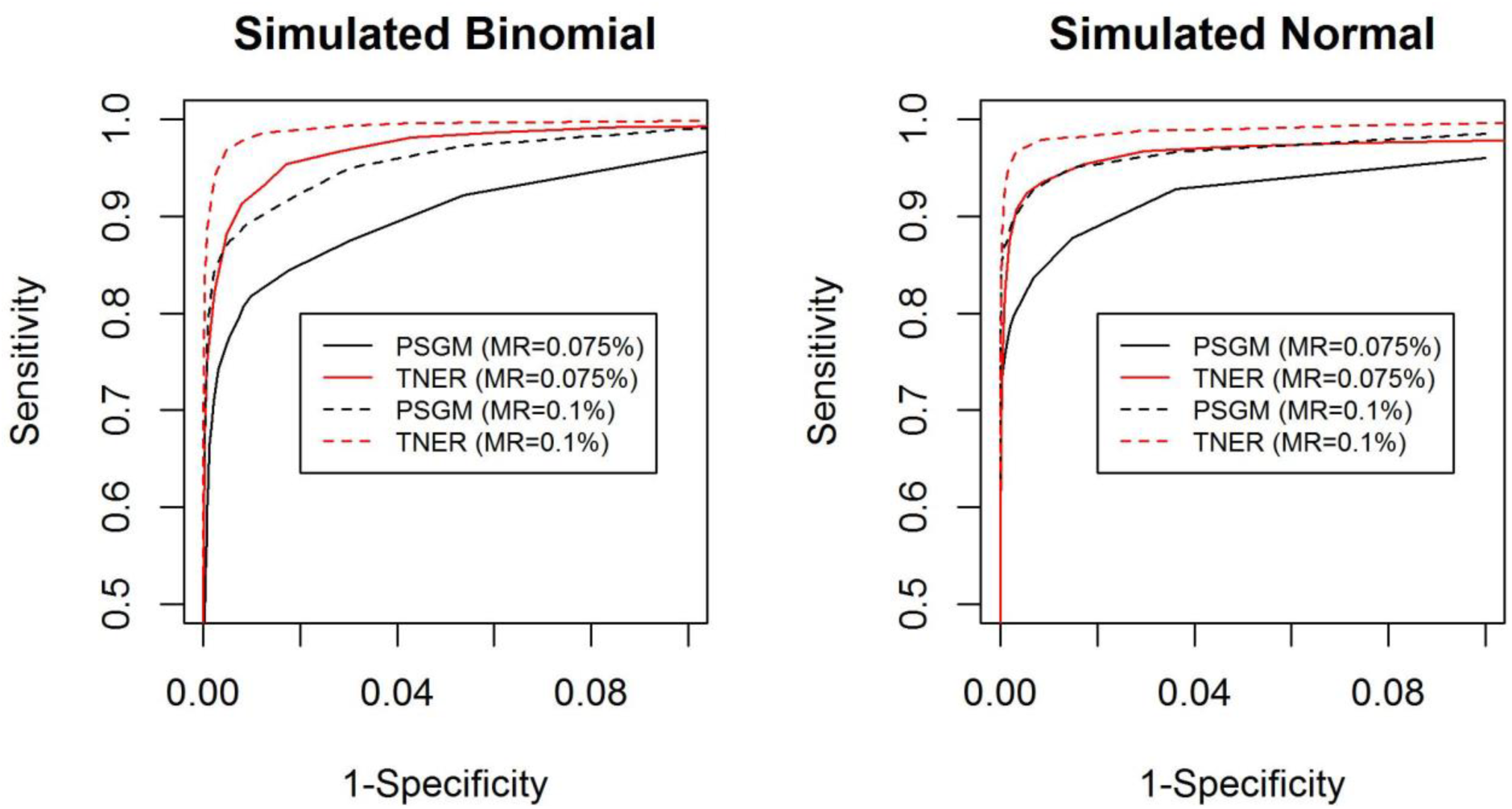
ROC curves for position specific Gaussian model (PSGM) (black) and TNER (red) methods in simulated cfDNA data. Two mutation rates (MR) were simulated: 0.075% (solid line) and 0.1% (dashed line) with a total coverage of 10000 at each position.

One of the advantages of the TNER method is that it uses information from other positions with the same TNC through a Bayesian consideration and thus stabilizes parameter estimates of the BMER. Therefore we would expect TNER performs better than position specific error models when the available sample size for healthy subjects becomes small. To evaluate the effect of healthy subject sample size on the performance of mutation detection methods, we used half the available healthy subjects (n=7) as our background mutation estimate and compared the results from both position specific Gaussian model and TNER in the simulation studies. As expected, we found that the smaller sample size of healthy subjects did not substantially reduce the performance of TNER, but greatly reduced the performance of the position specific Gaussian method (Figure 3), comparing to other methods. This clearly illustrates the robustness of the TNER method when the number of healthy subjects is small. In fact we found TNER can work even with 1-3 healthy subjects without sacrificing too much in performance.

**Figure 3.**
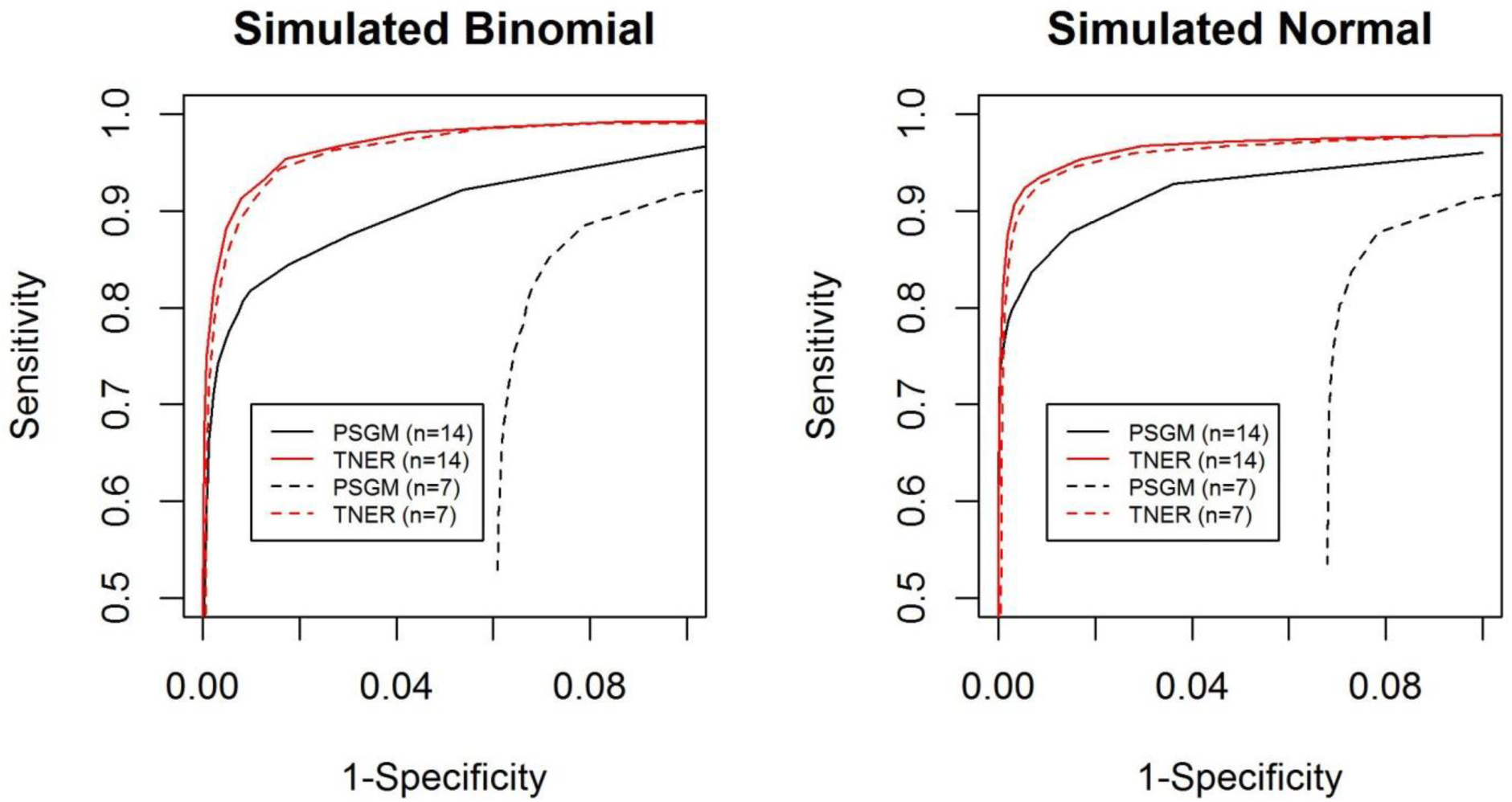
ROC curves of the position specific Gaussian model (PSGM) (black) and the TNER (red) methods with different input numbers of healthy subjects: n=7 (dashed line) and n=14 (solid line). Mutation rate used is 0.075%.

## 4. Discussion

In this study, we proposed TNER, a novel background polishing method for removing sequencing artifacts in panel sequencing data for liquid biopsy samples. The TNER method estimates background mutation errors from healthy subjects using a beta-binomial model to hierarchically incorporate both the tri-nucleotide level error rate and position specific error rate. The additional information from tri-nucleotide level data helps stabilize the estimate of background errors and proves to be more robust than the Gaussian based, position specific model used in iDES (Newman, et al., 2016), especially when the number of healthy subjects is small. Results on both simulated and real healthy subjects’ data demonstrated better performance of TNER than iDES in error reduction indicated by substantially more error-free positions and lower panel-wide error rate. TNER’s superior specificity in reducing false positive error can greatly benefit the down-streaming variant calling using general variant callers such as VarScan (Koboldt, et al., 2012) or MuTect (do Valle, et al., 2016).

We could have used dinucleotide context or more complicated local sequence context such as pentanucleotide (2 flanking nucleotides on each side) or heptanucleotide (3 flanking nucleotides on each side). The larger local sequence context may provide better model fit to the mutation error rate (Aggarwala and Voight, 2016), but the increasing model complexity with pentanucleotide (1,536 unique contexts) and heptanucleotide (24,576 unique contexts) becomes impractical for a targeted panel, like the one tested here with a total of 147k bases. The Bayesian prior parameter will not be well estimated due to small number of bases within each context. TNC provided a better fit than dinucleotide (Chen, et al., 2016) but not too complicated than the larger local sequence context (Aggarwala and Voight, 2016), so is a more balanced approach for a common NGS targeted panel.

One of the assumptions for analyzing NGS data by TNER is that individual nucleotides within a TNC share a more similar mutation error rate than those between TNC. We looked at the average mutation error rate from healthy subjects at the TNC level and compared the intra-TNC variability and the inter-TNC variability. About 94% of TNC have intra-TNC variability smaller than the inter-TNC variability. Figure 4 displays an example of three TNC, all with C to T substitution, showing very different distribution. The dashed lines are fit of beta distribution using the parameter estimates calculated by the method of moments. In general, beta distribution fits the intra-TNC error rate very well.

**Figure 4.**
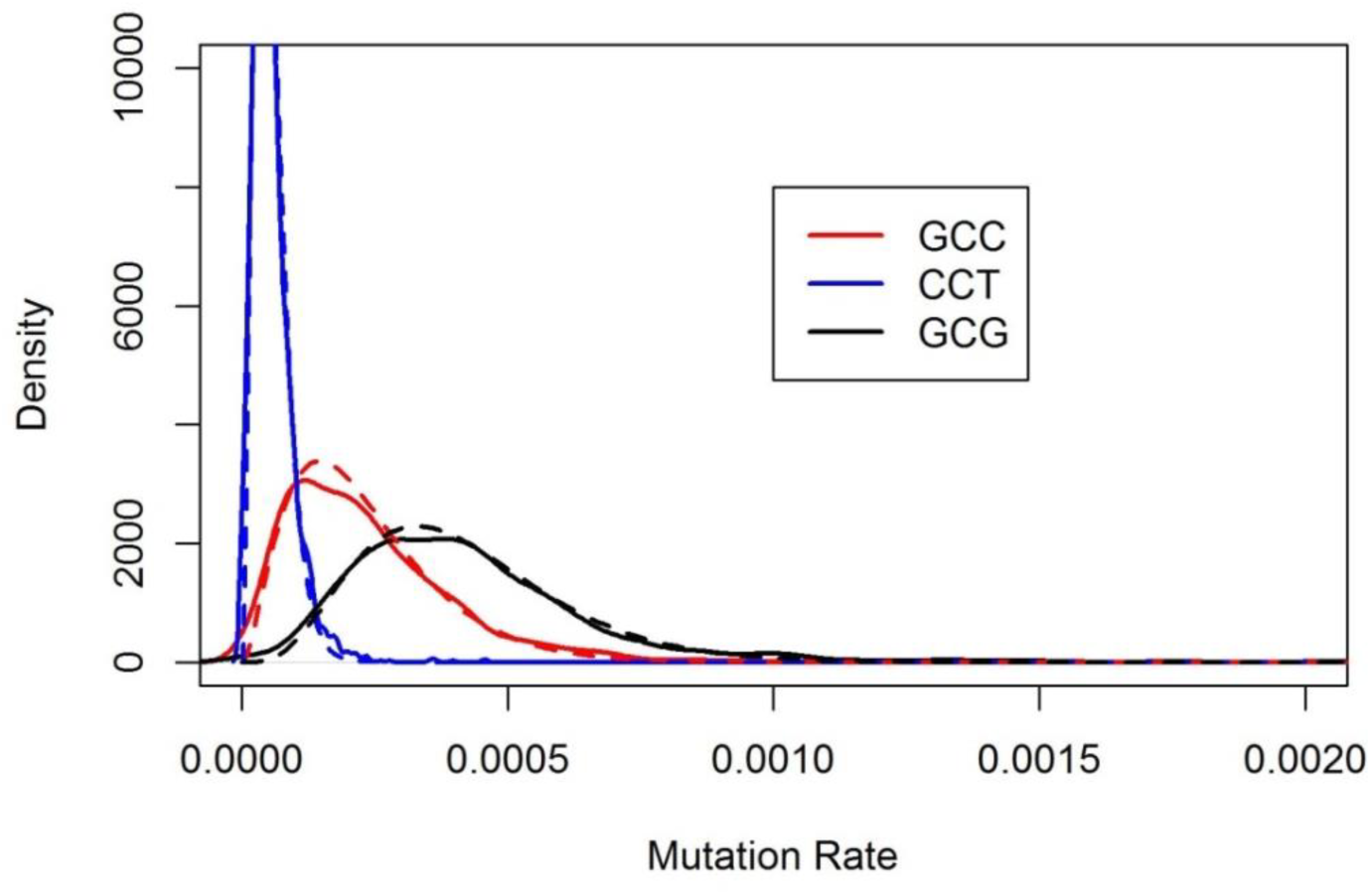
Examples of mutation error rate distribution of TNC with C-T substitution. Solid lines are the probability density of average position specific error rate within a TNC. The dashed lines are corresponding fit of a beta distribution using estimated parameters from the data.

In genomic data analysis, when the sample size is small, it is common to analyze data for individual genes using information from other genes. This is implemented in the *limma* method (Smyth, 2004) for microarray data analysis and the DESeq method (Anders and Huber, 2010) for RNAseq data analysis. In our approach, we take advantage of the large number of bases shared in the same nucleotide context and use these data to model the individual base mutation error rate. We found the TNER method improves the imprecise estimate of background associated with small sample size at individual base level.

Sequence data are read counts that are best described by distribution from discrete data families, such as Poisson distribution or binomial distribution, particularly when the read count is low and mutation frequency is very low such as in ctDNA data. We found that the Poisson distribution fit the count data well in general. A more sophisticated distributions taking over-dispersion into consideration and the zero-inflated nature of ctDNA data may further improve the method.

Currently, ctDNA is rapidly becoming established as an important tool to supplement conventional biopsies for cancer early detection, molecular characterization and monitoring of tumor dynamics. TNER method provides a novel approach to effectively reduce background noise in panel sequencing data for more accurate mutation detection in ctDNA.

---

**Supplementary Figure 1:**
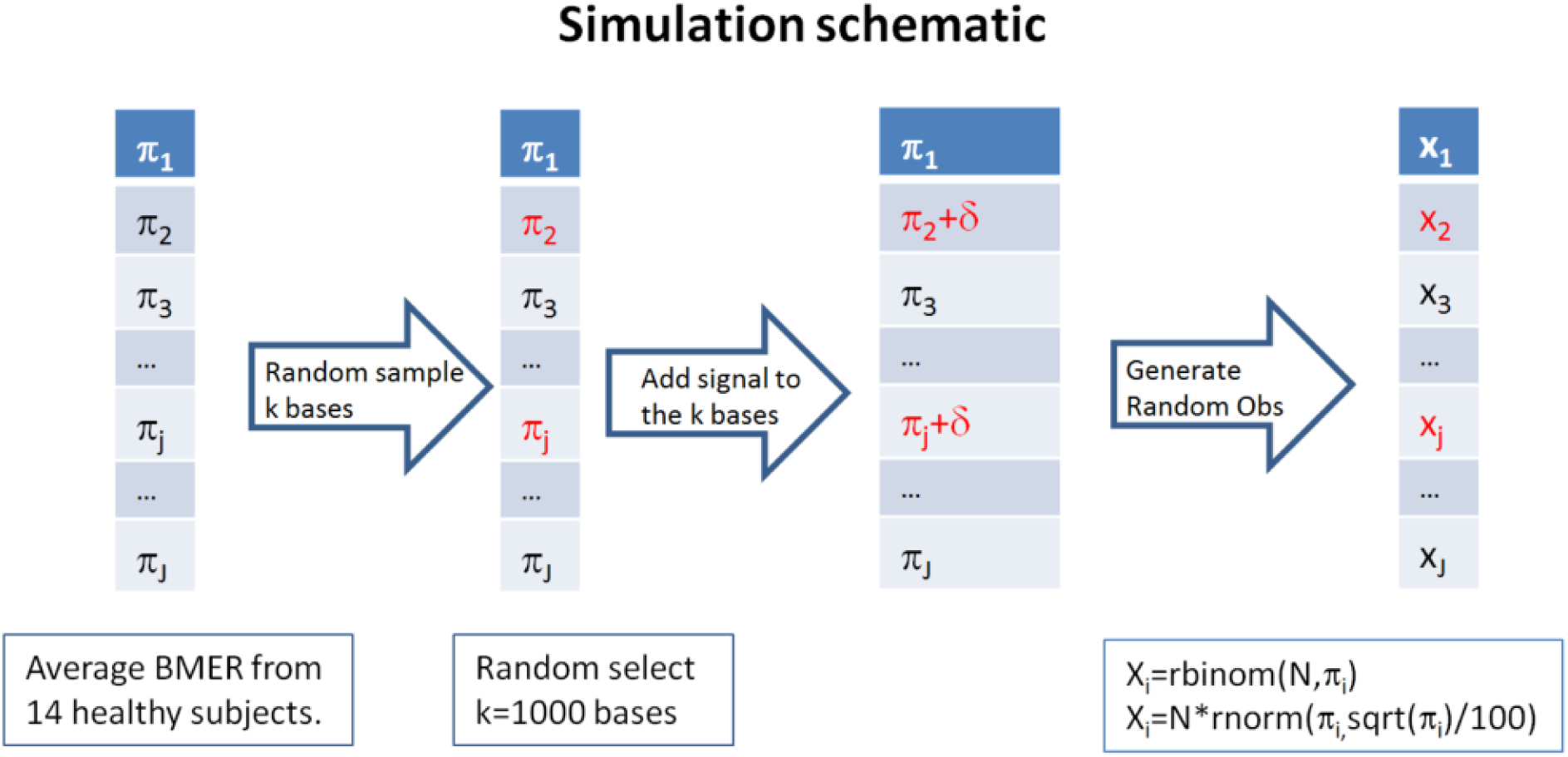
This illustrates how the simulated BMER (background mutation error rate) data were generated in one of four possible changes of a reference nucleotide.

